# Limited Dispersal of Benthic Environmental DNA from a Subtropical Mesophotic Shelf-Edge Bank

**DOI:** 10.1101/2024.08.26.609783

**Authors:** Luke J. McCartin, Annette F. Govindarajan, Jill M. McDermott, Santiago Herrera

## Abstract

Environmental DNA (eDNA) offers a powerful, non-invasive means of assessing biodiversity in marine ecosystems, yet the spatial resolution of eDNA remains poorly understood. We investigated the vertical and horizontal dispersion of eDNA from an isolated mesophotic coral reef (Bright Bank) in the stratified offshore waters of the northern Gulf of Mexico’s shelf edge. We conducted comprehensive vertical and horizontal water column eDNA sampling across multiple radial directions and depths. We characterized invertebrate communities using a paired metabarcoding approach targeting broad (18S) and taxon-specific (28S) markers. We found that vertical transport of benthic eDNA was limited by water column stratification, with distinct benthic community signals confined to the near-bottom layers. In contrast, horizontal dispersal of eDNA extended beyond at least 1.5 km, though the prevalence of eDNA from benthic invertebrates declined with increasing distance from the bank. Taxon-specific primers showed greater detection sensitivity and dispersal range, particularly for benthic corals, than primers that are used to broadly assess eukaryotic biodiversity. These findings demonstrate that water column structure and marker selection critically influence the spatial interpretation of marine eDNA data. The study represents a snapshot of late-summer conditions. Seasonal variability should be considered in future studies. Our results provide a realistic framework for integrating eDNA into offshore environmental surveillance, biodiversity monitoring, and spatial management.

## Introduction

Marine biodiversity underpins ecosystem function and resilience, yet its accurate assessment remains challenging, particularly in offshore and remote marine environments. The rapid development of environmental DNA (eDNA) approaches, where DNA extracted from seawater is used to detect species presence and assess biodiversity, has significantly advanced marine biodiversity monitoring capabilities (Patin and Goodwin, 2023; Yang et al., 2024). These molecular methods complement traditional sampling and visual surveys, often providing broader, more sensitive species detection (Govindarajan et al., 2021; Govindarajan et al., 2023; West et al., 2024). Despite their advantages, accurately interpreting the source of detected eDNA signals in marine ecosystems remains challenging. Once released from organisms, marine eDNA disperses vertically and horizontally through physical processes such as dilution, diffusion, and advection by currents, before settling to the benthos or degrading into undetectable shorter fragments (Harrison et al., 2019). Therefore, accurately inferring species presence and location from eDNA relies fundamentally on understanding how eDNA disperses—a core factor governing the "ecology of eDNA" (Barnes & Turner, 2016).

Insights into marine eDNA dispersion dynamics primarily stem from two complementary approaches: (1) hydrodynamic modeling using particle-tracking simulations and (2) empirical eDNA field sampling around known eDNA sources. Both approaches consistently indicate that physical and ecological factors can significantly constrain vertical dispersion. For instance, pronounced stratification, such as haloclines common in fjords, creates distinct vertical eDNA differences, even at small, meter-scale depth intervals (Jeunen et al., 2020; Robinson et al., 2023). Similarly, distinct vertical eDNA patterns have been observed in structurally complex habitats such as kelp forests, with clear differentiation between surface and subsurface communities (Monuki et al., 2021). Limited vertical eDNA dispersion is also apparent in the ocean’s interior, where different water masses harbor unique eDNA signatures (Govindarajan et al., 2021; Canals et al., 2021). These field observations are consistent with modeling results (Allan et al., 2021). In contrast, regions characterized by active vertical mixing, such as upwelling zones, display minimal vertical differentiation in eDNA signatures (Closek et al., 2019).

While modeling studies predict potential horizontal eDNA transport distances extending tens of kilometers (Andruszkiewicz et al., 2019; Kutti et al., 2020), empirical field studies often report localized, patchy eDNA detections even in well-connected marine systems (Port et al., 2016; Jeunen et al., 2019; West et al., 2020; Dugal et al., 2023; Shea & Boehm, 2024). This discrepancy arises because physical transport models frequently omit critical ecological and molecular processes such as sedimentation, adsorption onto particulate matter, and varying amplification efficiencies that determine detection limits and persistence (Harrison et al., 2019; Kelly et al., 2019; Collins et al., 2018). Therefore, empirical studies with spatially explicit sampling designs are critical to validate model predictions, refine our understanding of eDNA dispersal distances, and enhance the ecological interpretation of eDNA data.

A knowledge gap exists because empirical tests of generalizable hypotheses about marine eDNA dispersion have rarely been conducted in offshore marine ecosystems. Even fewer studies use clearly identifiable, isolated natural eDNA sources. Two central hypotheses exist: (1) the vertical dispersion of benthic-derived eDNA is constrained primarily by water column stratification, and (2) eDNA detection decreases rapidly with increasing horizontal distance from its source. Empirical validation of these hypotheses in offshore contexts remains limited (but see Baetscher et al. 2024; Shea et al. 2022; Ely et al., 2021; Murakami et al., 2019), restricting our ability to accurately interpret eDNA signals in these environments.

This study aims to fill this critical gap in existing knowledge by empirically testing these hypotheses at Bright Bankan isolated mesophotic reef in the northwestern Gulf of Mexico that is uniquely well suited as a natural laboratory for spatial eDNA dispersion studies. Leveraging an isolated benthic invertebrate community as a clearly defined eDNA point source, we systematically collected seawater samples at multiple depths and horizontal distances from the bank. Additionally, by comparing results derived from metabarcoding with universal eukaryotic (18S rDNA) primers and anthozoan coral-specific (28S rDNA) primers, we evaluated how primer specificity influences the spatial interpretation of marine eDNA signals. Findings from this research will significantly improve the interpretation and enhance the precision of eDNA-based marine biodiversity assessments, optimize field sampling strategies, and inform conservation and resource management decisions in offshore marine ecosystems.

## Materials and Methods

### 2.1 Study Site

The northwestern Gulf of Mexico contains steep-sloped carbonate banks formed by salt diapirs along the continental shelf edge, supporting distinct benthic invertebrate communities, including corals and sponges (Rezak et al., 1985; Dennis & Bright, 1988). Seasonal temperature fluctuations lead to pronounced summer stratification, creating distinct ecological zones at different depths (Lugo-Fernández, 1998; Sammarco et al., 2016). Bright Bank, within the Flower Garden Banks National Marine Sanctuary (FGBNMS), lies approximately 108 nautical miles (NM) south of the Louisiana-Texas coast, rising from ∼120 m at its base to a peak of ∼33 m below sea level. The bank’s upper surface predominantly ranges between 40 and 80 m in depth and is isolated from nearby hard substrates such as Rankin & 28 Fathom Banks (∼8 NM west) and Geyer Bank (∼12 NM east) (**Figure 1A**). The isolation of Bright Bank’s benthic community makes it ideal for studying spatial patterns of eDNA dispersion.

**Figure 1.**
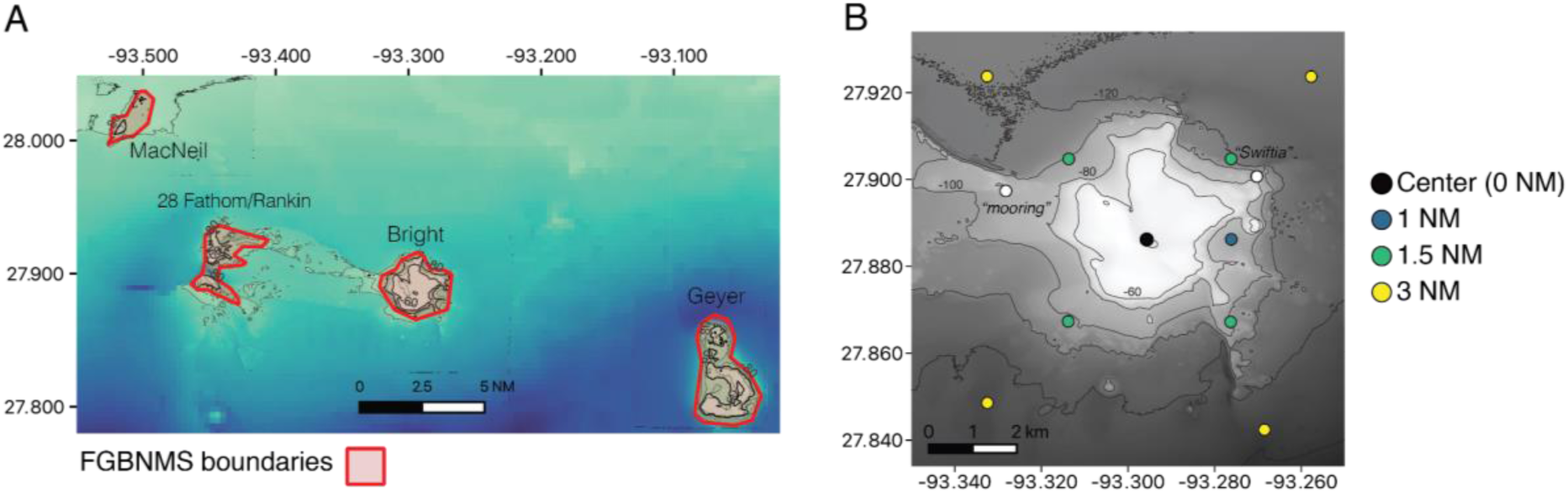
Study site overview and sampling station locations at Bright Bank. A) Bathymetric map of the study region in the northwestern Gulf of Mexico. Bathymetry is from the United States Geological Survey (USGS) (Gardner and Beaudoin, 2005) and the General Bathymetric Chart of the Oceans (GEBCO Bathymetric Compilation Group, 2023). B) Zoomed-in bathymetry of Bright Bank with 20-meter contour intervals. Station colors represent distances from the bank’s center, except for casts at the “*Swiftia”* and “mooring” stations.

### 2.2 Study Design

We conducted an observational study from September 24–26, 2019, aboard *R/V Manta*, employing systematic spatial sampling to test hypotheses related to eDNA vertical and horizontal dispersion. Sampling stations included the bank’s center and radial points at distances of 1.0, 1.5, and 3.0 NM **(Figure 1B**). Distances and sampling depths were selected to capture the anticipated scale of eDNA dispersion. Two additional sites (“mooring” and “*Swiftia*”) were sampled ∼2 NM north of Bright Bank, corresponding to previously explored remotely operated vehicle (ROV) dive sites. Sampling near the seafloor at these sites was conducted to groundtruth the detection of eDNA from corals known from these sites at depths of ∼80 meters. Sampling occurred during daylight hours between 07:00 and 19:05 local time (CST) (**Table 1, Table S1**).

**Table 1:** Coordinates of CTD cast stations and eDNA sampling depths.

| Cast | Latitude | Longitude | Station | Depths sampled (Niskin replicates) |
| --- | --- | --- | --- | --- |
| 1 | 27.88619 | -93.29568 | Center | 20 m (4), 40 m (4), 58 m (4) |
| 3 | 27.90475 | -93.313693 | NW - 1.5 NM | 40 m (3), 60 m (3), 80 m (3), 95 m (3) |
| 4 | 27.92374 | -93.33255 | NW - 3.0 NM | 40 m (3), 60 m (3), 80 m (3), 100 m (2) |
| 5 | 27.90476 | -93.27621 | NE - 1.5 NM | 40 m (3), 60 m (3), 81 m (3), 101 m (3) |
| 8 | 27.84239 | -93.268503 | SE - 3.0 NM | 40 m (3), 60 m (3), 80 m (3), 100 m (3) |
| 9 | 27.86722 | -93.27632 | SE - 1.5 NM | 40 m (3), 60 m (3), 80 m (3), 96 m (3) |
| 10 | 27.86735 | -93.31381 | SW - 1.5 NM | 40 m (3), 60 m (3), 80 m (3), 100 m (3) |
| 12 | 27.923706 | -93.257714 | NE - 3.0 NM | 40 m (3), 60 m (3), 80 m (3), 100 m (2) |
| 13 | 27.84857 | -93.33247 | SW - 3.0 NM | 40 m (3), 60 m (3), 80 m (3), 100 m (3) |
| 16 | 27.88623 | -93.27616 | E - 1.0NM | 40 m (3), 60 m (3), 80 m (3), 90 m (3) |
| 2 | 27.89738 | -93.32821 | "Swiftia" | 84 m (all Niskins over one filter) |
| 11 | 27.900727 | -93.270305 | "mooring" | 83 m (all Niskins over one filter) |

Water samples were collected using a Niskin bottle rosette at depths from 20 to 100 meters below sea level at 20-meter intervals. At locations with bottom depths shallower than 100 meters, we collected samples as close to the seafloor as possible based on bathymetry data - within 6 m of the bottom. At the bank’s center, four replicate 2L samples were independently collected at each depth (i.e. separate Niskin bottles), while three replicate samples per depth were collected at other stations. Samples with an average volume of 2.2 ± 0.21 L (1 SD) were immediately filtered after instrument recovery through separate 0.22 µm polyethersulfone (PES) Sterivex filters (Millipore) using peristaltic pumps (Govindarajan et al., 2021). At “Mooring” and “*Swiftia*” stations, water was sampled near the seafloor, filtering the entire volume of all Niskin bottles (28.2 and 30 L, respectively) through 0.20 µm PES Mini Kleenpak capsules (Pall). Negative controls of MilliQ water were filtered alongside field samples (Sterivex average volume = ∼2.37 ± 0.14 L; Mini Kleenpak average volume = 22 L) to monitor potential contamination. All filtration followed stringent contamination prevention protocols as detailed in Govindarajan et al. (2022).

### 2.3 Environmental Profiling

Conductivity, temperature, depth (CTD), fluorescence, turbidity, and dissolved oxygen were measured concurrently with water sampling. Measurements were made using a Sea-Bird SBE 19plus V2 CTD mounted on the rosette, as well as a WET Labs ECO-AFL/FL fluorescence sensor, a WET Labs ECO turbidity sensor, and a SBE 43 dissolved oxygen sensor (Seabird Scientific, Bellevue, WA). Data from downcasts were analyzed to generate environmental profiles.

### 2.4 DNA Extraction and Metabarcoding

DNA was extracted using DNEasy Blood & Tissue Kits (Qiagen) with modified protocols optimized for Sterivex and Kleenpak filters (Govindarajan et al., 2021; Govindarajan et al., 2022). Metabarcoding employed two primer sets: broad-range 18S rDNA primers targeting diverse eukaryotes (Amaral-Zettler et al., 2009) and a taxon-specific 28S rDNA set focused on anthozoan corals and related invertebrates (McCartin et al., 2024). Combining these markers provided both broad biodiversity assessments and focused insights into coral communities. Protocols for amplification using these primers are provided in Govindarajan et al. (2022) (18S) and McCartin et al. (2024) (28S). Differences between these protocols are summarized in **Table 2**. Following amplification and tagging, PCR products were pooled and sequenced using Illumina MiniSeq and MiSeq platforms, including sampling and PCR negative controls (molecular grade water).

**Table 2:**
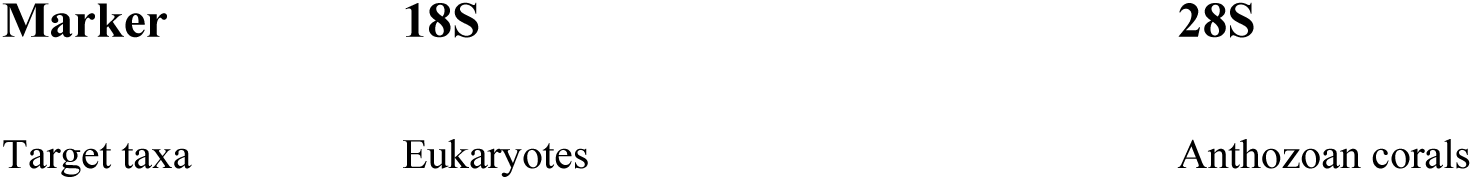
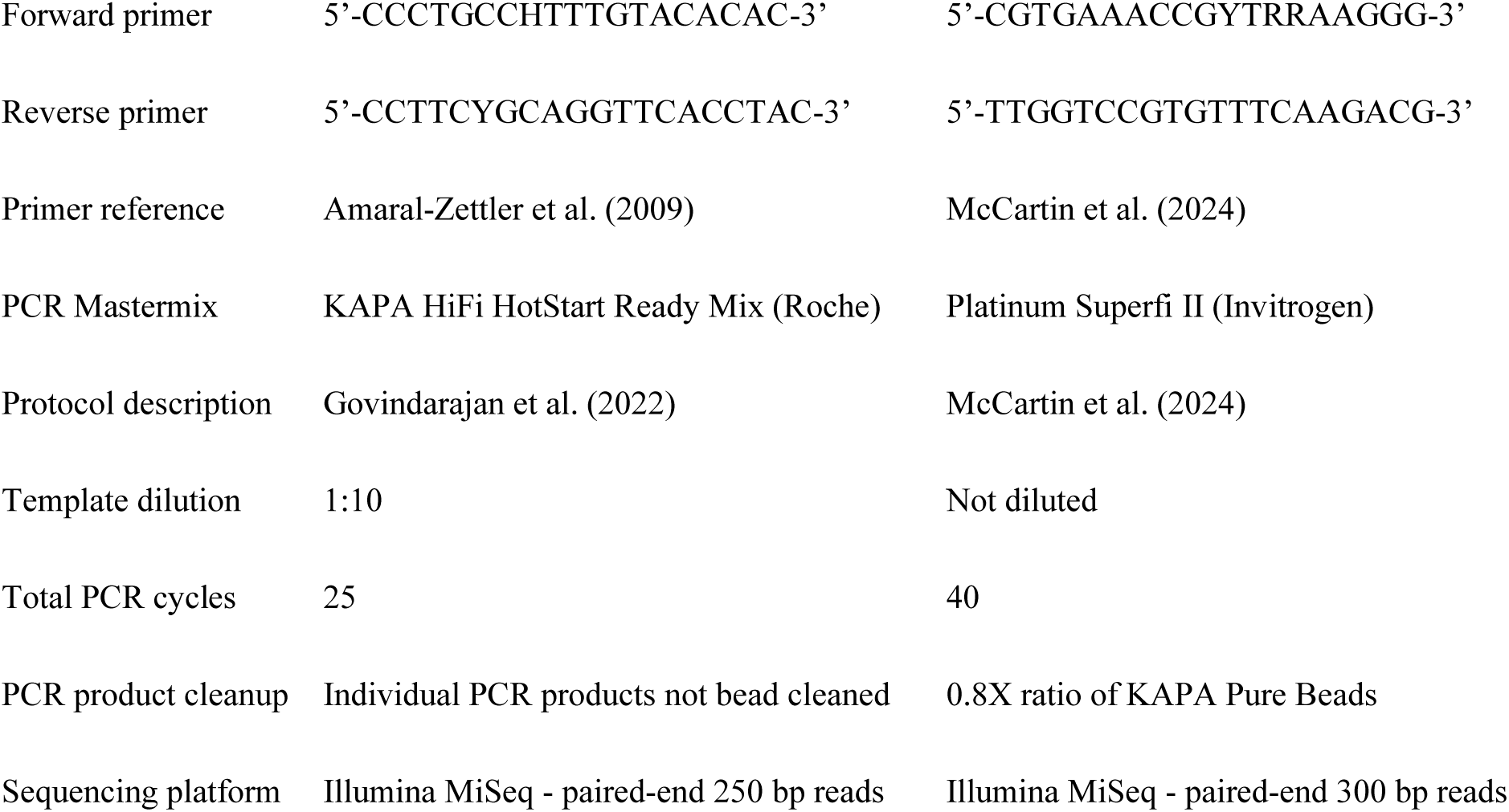
Differences between PCR amplification protocols for metabarcoding 18S and 28S markers.

### 2.5 Bioinformatic Processing

Raw sequencing reads underwent trimming, quality filtering, denoising, and chimera removal using *cutadapt* and *DADA2* (Martin, 2011; Callahan et al., 2016). Amplicon Sequence Variants (ASVs) were classified taxonomically using a naïve Bayesian classifier implemented in *DADA2* (Wang et al., 2007), referencing custom databases compiled from MetaZooGene and curated anthozoan sequences (O’Brien et al., 2024; McCartin et al., 2024). For 18S, global data for all taxa were downloaded from MetaZooGene on April 25, 2024. For *28S*, data from genera and/or species known from the North Atlantic were downloaded from MetaZooGene on June 20, 2024. Following taxonomic classification, ASVs identified as contaminants from sampling and PCR negative controls were removed using the R package *decontam* (Davis et al., 2018). A detailed description of the bioinformatic analyses are provided in Appendix A: Supplementary Methods.

### 2.6 Data Analysis

Data analysis and visualization were conducted in *R* (version 4.3.1) using *phyloseq* (McMurdie & Holmes, 2013), *vegan* (Oksanen et al., 2007) and *tidyverse* (Wickham et al., 2019) packages. Rarefaction curves assessed sequencing completeness. The data were subset to contain ASVs classified to animal phyla. This animal-only subset was used to explore biodiversity patterns described in this paper. Bray-Curtis dissimilarities were calculated based on the presence/absence (Clarke et al., 2014) of ASVs across sampling stations and depths. Distance matrices underwent Principal Coordinates Analysis (PCoA) and distance-based redundancy analysis (dbRDA) to test for associations with depth, spatial coordinates, and proximity to the seafloor. Relationships between PCoA axes and depth were investigated using linear regression.

Marine invertebrate families were classified as benthic based on established ecological life-history traits (Brusca et al., 2022; Jumars et al., 2015). Families producing pelagic larvae or with pelagic life stages (e.g. sessile, colonial hydroids) but with predominantly benthic adult forms were classified as benthic for this analysis. We used Analysis of Compositions of Microbiomes with Bias Correction 2 (ANCOM-BC2) (Lin & Peddada, 2024) to determine if eDNA from benthic invertebrates was differentially abundant across sample replicates taken at different stations and depths.

Bathymetric maps were created using *QGIS*, *marmap* (Pante & Simon-Bouhet, 2013) with bathymetric data from USGS and GEBCO (Gardner et al., 1998; GEBCO Compilation Group, 2023).

### 2.7 Analysis and Correction for Potential Contamination

Significant contamination detected in negative controls associated with 18S data from one cast (Cast 16 - Station 1.0 NM East) led to the exclusion of this data from the analysis. Minor contamination detected in some negative controls was assessed and determined to be insufficient to influence overall interpretations; therefore, no further data adjustments were applied. A detailed summary of ASVs detected in negative controls and corrective measures are provided in Appendix A: Supplementary Methods, and comprehensive read counts and taxonomic classifications for all ASVs across all samples are available in **Tables S2** and **S3**.

## 3. Results

### 3.1 Water column profile

A strong and consistent thermocline formed below the surface mixed layer, beginning at depths of 40–50 meters during the entire sampling period (**Figure 2**). This thermocline is typical for the northwestern Gulf of Mexico in late summer (Lugo-Fernández, 1998). This thermocline coincided with reductions in salinity and dissolved oxygen and increases in fluorescence and turbidity, clearly marking the chlorophyll maximum zone at this transition (**Figure S1**).

**Figure 2:**
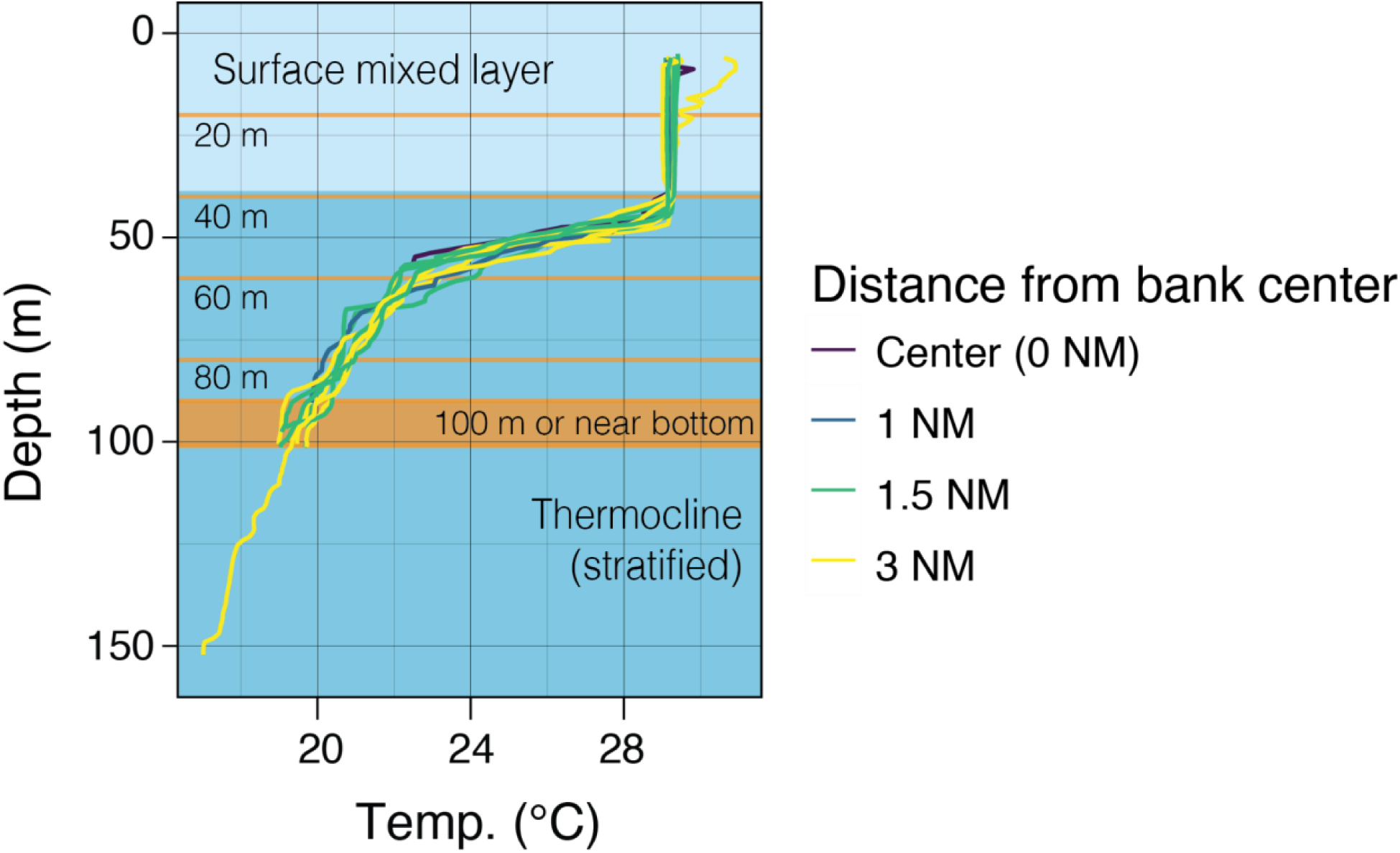
Temperature profiles measured during CTD casts conducted for eDNA sampling over the center and at radial stations distances away from Bright Bank. Sampling depths are indicated in orange and labeled for reference.

### 3.2 Amplicon sequence diversity and sequencing depth

Post-processing yielded an average sequencing depth of 118,676 ± 26,704 reads per sample for the 18S dataset and 45,971 ± 15,571 reads per sample for the 28S dataset. The average ASV richness per sample was substantially higher for the 18S dataset (891 ± 217 ASVs) compared to the 28S dataset (254 ± 70 ASVs). Rarefaction analyses indicated sufficient sequencing coverage for both markers, with more pronounced saturation in the 18S dataset (**Figure S2**).

### 3.3 Taxonomic composition of eDNA

Sequencing reads classified to animal phyla (hereafter animal reads) represented 43.5 ± 20.7% of the 18S dataset and 68.0 ± 19.6% of the 28S dataset. Animal ASVs accounted for a larger proportion of the 28S dataset (25.2 ± 9.3%) compared to the 18S dataset (5.2 ± 1.4%). In the 18S dataset, copepods (classes Calanoida and Cyclopoida, phylum Arthropoda) dominated animal reads by percentage (86.8 ± 14.4%; **Figure S3**). Hydrozoans (phylum Cnidaria, primarily siphonophores) comprised 9.0 ± 13.9% of animal reads. Additional taxa that comprised at least 5% of animal reads in any sample included Appendicularia, Thaliacea (Chordata); Phascolosomatidea, Polychaeta (Annelida); Chaetognatha; Tentaculata (Ctenophora); and Demospongiae (Porifera). Mollusca, Bryozoa, Echinodermata, Platyhelminthes, Nemertea, and Rotifera were present at low abundances (<5% in every sample). In the 28S dataset, hydrozoans dominated animal reads (78.7 ± 21.2%), followed by Tentaculata (Ctenophora, 13.0 ± 16.3%). Anthozoans and calcareous sponges each comprised >10% of animal reads in multiple samples. Other groups present in the 28S data included Nuda (Ctenophora), Scyphozoa (Cnidaria), Cephalopoda (Mollusca), Annelida, Chordata, and Arthropoda.

### 3.4 Vertical Distribution of Invertebrate eDNA

Principal Coordinates Analysis (PCoA) based on Bray–Curtis dissimilarities distinguished surface mixed layer samples (20 and 40 m depths) clearly from deeper samples (≥ 60 m; **Figure 3**). The first PCoA axes significantly inversely correlated with depth (*18S* P < 0.00001, Adjusted R^2^ = 0.7008; *28S* P < 0.00001, Adjusted R^2^ = 0.8612), and explained 22.7% of the variation in 18S data and 14.4% in 28S data. Distance-based redundancy analysis (dbRDA) further supported these findings, constraining 27.7% (*18S*) and 21.6% (*28S*) of community variation. Bray–Curtis dissimilarities were significantly correlated with depth in both datasets ([18S P = 0.001, 28S P = 0.001]; **Tables 3 and 4**).

**Figure 3.**
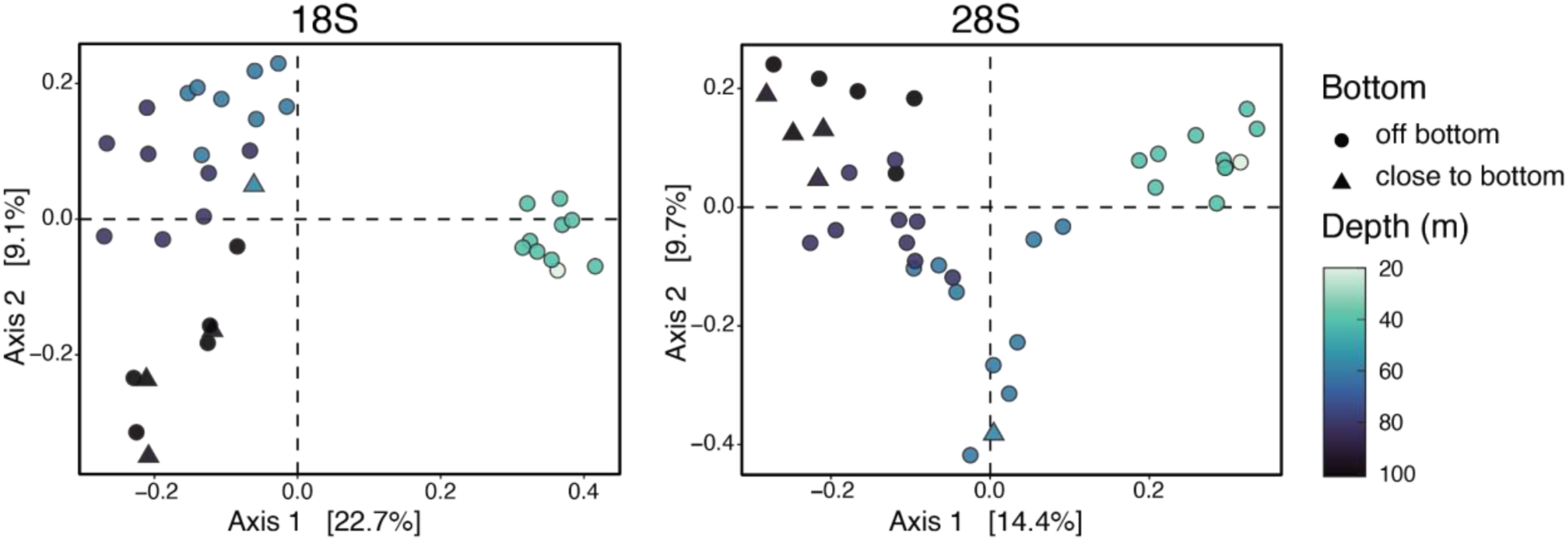
Multivariate community similarity of eDNA samples across depth and markers. Principal coordinates analysis (PCoA) plots based on Bray–Curtis dissimilarities for *18S* and *28S* marker datasets. Points represent aggregated replicate data per station and depth.

**Table 3:** Results of permutational ANOVA to test for significance in distance-based redundancy analyses of 18S data.

| Term | Degrees of Freedom | Sum of Squares | F | Pr(>F) |
| --- | --- | --- | --- | --- |
| Depth | 1 | 1.5712 | 7.4257 | 0.001*** |
| Latitude | 1 | 0.2195 | 1.0375 | 0.331 |
| Longitude | 1 | 0.2505 | 1.1841 | 0.204 |
| Proximity to bottom (Y/N) | 1 | 0.4004 | 1.8924 | 0.011* |
| Residual | 30 | 6.3476 |  |  |

**Table 4:** Results of permutational ANOVA to test for significance in distance-based redundancy analyses of 28S data.

| **Term** | **Degrees of Freedom** | **Sum Of Squares** | **F** | **Pr(>F)** |
| --- | --- | --- | --- | --- |
| Depth | 1 | 1.3074 | 5.5840 | 0.001*** |
| Latitude | 1 | 0.3464 | 1.4793 | 0.034* |
| Longitude | 1 | 0.2397 | 1.0239 | 0.357 |
| Proximity to bottom (Y/N) | 1 | 0.3007 | 1.2844 | 0.110 |
| Residual | 34 | 7.9605 |  |  |

### 3.5 Dispersion of eDNA from Benthic Invertebrates

The richness of benthic invertebrate families peaked near the seafloor at the bank’s center (27.3 ± 4.9 families for *18S*; 15.8 ± 0.5 families for *28S*) and declined substantially at 60 m depth with increasing horizontal distance away (**Figure 4**). Coral eDNA genera richness showed similar spatial patterns in the *28S* data, consistently declining with distance. Likewise, the abundance of eDNA from several benthic invertebrate families, including corals, was significantly correlated with depth and distance from Bright Bank’s center. In the 18S data, the abundance of eDNA from benthic invertebrates, including corals, sponges, polychaetes, and bryozoans, was significantly lower with increasing distance from the center of Bright (**Table S4**). In the 28S data, the abundance of eDNA from sponges, corals, and sessile hydroids was significantly lower with increasing distance from Bright (**Table S5**). Notably, eDNA from some coral families (e.g., Aphanipathidae) was positively correlated with depth but not with distance from the center of Bright.

**Figure 4:**
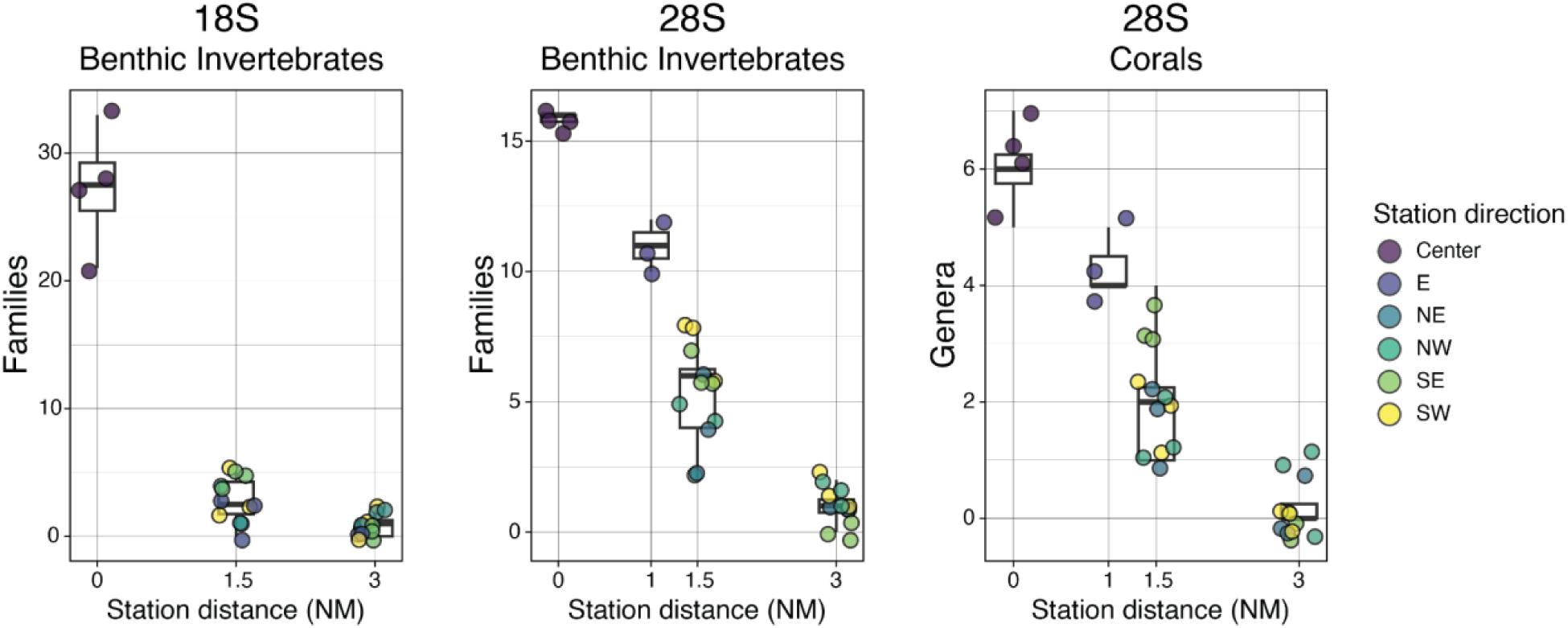
Horizontal patterns in benthic eDNA richness at 60 m depth. Richness of benthic invertebrate taxa at 60 m depth at sampling stations varying distances from the center of Bright Bank. Each point represents a replicate sample.

eDNA from corals was consistently and inversely correlated with distance from Bright Bank across both datasets and both differential abundance analyses; however, coral eDNA was detected at substantial distances away in the 28S data. Black coral eDNA and eDNA from octocorals were detected in all sample replicates as far as 1.5 NM (∼2.7 km) away from the center of Bright Bank, particularly at 60 and 80 m at eastern, southeastern, and southwestern stations (**Figure 5A**). On the contrary, coral eDNA was not consistently detected across sample replicates in the 18S data at any site other than the center of Bright Bank (**Figure 5B**). Overall, the relative abundance of coral eDNA was substantially less in the 18S data; at maximum, 0.2% of the animal eDNA in any 2L eDNA sample and 3% in the large-volume samples. The 28S libraries were comparatively enriched in eDNA from corals; at maximum, 24.6% in the 2L samples and nearly 50% in the large volume samples.

**Figure 5:**
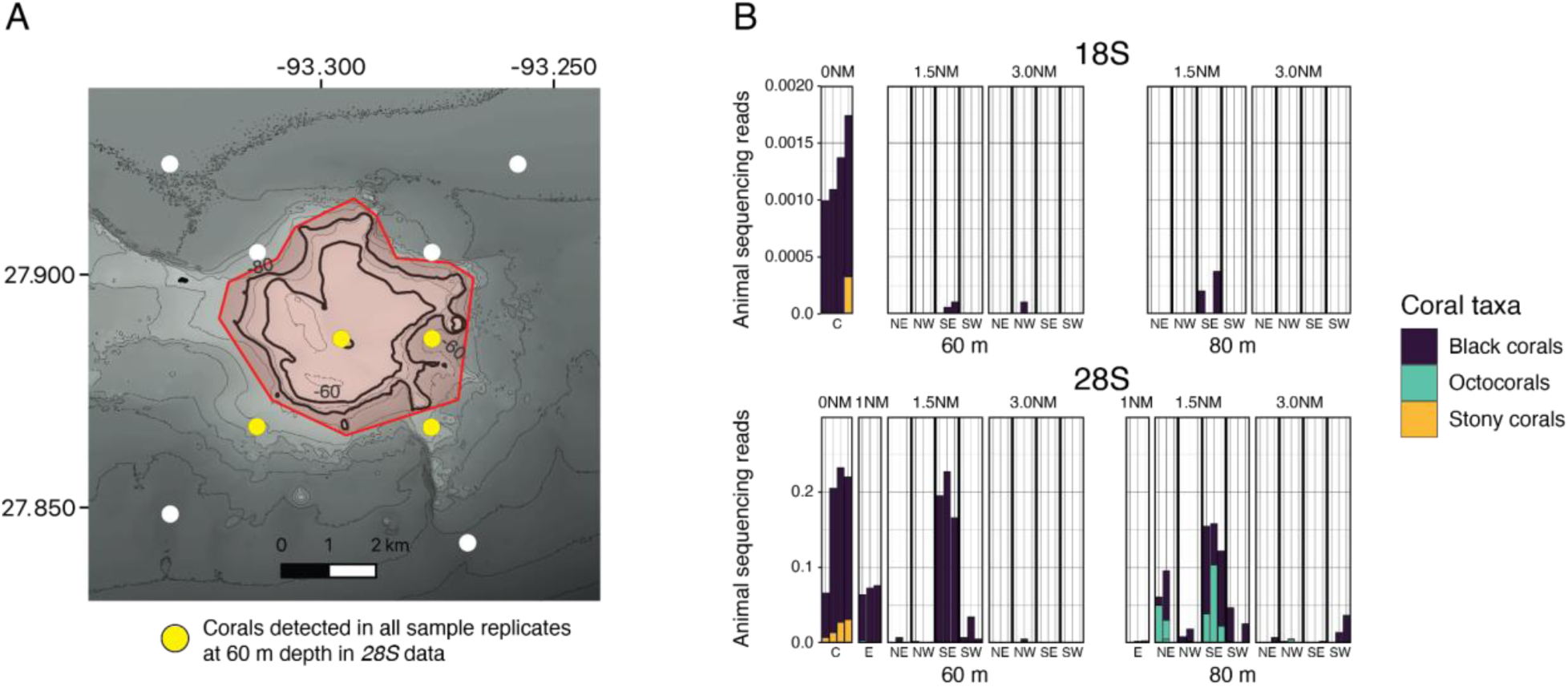
Spatial detection of coral eDNA across depths and directions from Bright Bank. **A)** Presence of coral eDNA at 60 and 80 m depths across radial transects. Yellow points indicate high-confidence detections (all replicates positive). Isobath contours (60 m and 80 m) and the boundary of the Flower Garden Banks National Marine Sanctuary (FGBNMS) are indicated. **B)** Proportion of animal sequencing reads classified as coral across sequencing datasets, sampling stations, and depths of 60 and 80 meters.

## 4. Discussion

### 4.1 Vertical eDNA Transport is Limited by Water Column Stratification

Our findings clearly support the hypothesis that vertical eDNA transport is constrained by water column stratification, consistent with prior studies (Jeunen et al., 2019; Allan et al., 2021; Govindarajan et al., 2021; Monuki et al., 2021; Hoban et al., 2023; Robinson et al., 2023). Both metabarcoding datasets (18S and 28S) revealed significant vertical differentiation in animal eDNA communities. These data also revealed an increase in benthic-derived eDNA with depth for some groups of octocorals and black corals. These results align closely with direct observations at deeper habitats on the slopes of Bright Bank and to its east (Herrera & McCartin 2024). Such stratification-driven differences underscore that accurately assessing biodiversity through eDNA must account for variability in vertical mixing.

Detection of benthic invertebrate eDNA at depths in the surface mixed layer was rare (**Figure S4**), likely reflecting episodic physical or biological processes such as spawning events, vertical migrations, larval dispersal, or storm-induced mixing (Harrison et al., 2019). For example, the 18S dataset revealed notable detections of an opheliid polychaete in the surface mixed layer, comprising up to 10% of sequencing reads at 40 meters depth, potentially reflecting a reproductive migration event (Maciolek & Blake, 2006, Pandian 2019). Such episodic detections highlight the importance of integrating ecological and environmental context into eDNA sampling strategies, necessitating repeated temporal and spatial sampling to differentiate between persistent signals and transient biological events (Stommel 1963).

### 4.2 Horizontal transport of eDNA exceeds one kilometer

Benthic eDNA richness peaked near the seafloor at Bright Bank’s center (∼58 m depth), reflecting a diverse assemblage of corals, sponges, polychaetes, bryozoans, ascidians, and mollusks. Consistent with our hypotheses, both the richness and relative abundance of eDNA from benthic invertebrates rapidly declined with horizontal distance due to mixing, diffusion, and degradation processes. These results are consistent with a substantial number of studies that have found that eDNA metabarcoding distinguishes differences in biological communities that are physically connected by currents distances over similar or closer distances apart than our field stations (Port et al., 2016; Jeunen et al., 2019; West et al., 2020; Dugal et al., 2023).

While the diversity of benthic invertebrate eDNA decreased with distance from Bright Bank, eDNA transport was still apparent in these data. In the 28S data we consistently detected coral eDNA, particularly from black corals (Antipathidae) and ellisellid octocorals (*Ellisella* spp. and *Nicella* spp.), at stations up to approximately 2.8 kilometers (1.5 NM) from the bank center. Based on bathymetry data, these stations were up to 1.46 km from the seafloor at the sampling depths where corals were consistently detected (60 and 80 meters). The consistent detection of eDNA likely originating ∼1.5 kilometers from its source is similar to empirically measured transport distances of eDNA originating from caged fish in inlets and bays of marine eDNA. These studies found that eDNA may be dispersed at detectable concentrations at distances from 1 to 5 kilometers away (Murakami et al., 2019; Shea et al., 2022; Baetscher et al., 2024). Our study differs from these previous studies in that we measured eDNA dispersal at an offshore habitat where the coastline does not influence transport. Our study is also methodologically different in that we used metabarcoding, whereas these previous studies have used quantitative PCR (qPCR). Quantitative approaches like qPCR or digital droplet PCR that are complementary to metabarcoding should be employed in future studies to validate observed dispersal distances and provide absolute eDNA quantification. Even so, it is worth noting that linking eDNA quantification to organismal abundance and biomass is an area of active research with inherent uncertainties (Yates et al., 2025; Yates et al., 2019).

Studies incorporating models of ocean circulation and eDNA degradation have predicted the transport of detectable concentrations of eDNA over tens of kilometers (Andruskiewicz et al., 2019). Unfortunately, a circulation model that accurately hindcasts the current field at Bright Bank at a useful resolution (100 m or less) is not available. Generating such a model is a computationally intensive problem beyond the scope of this study. Still, we can use a model of eDNA persistence (McCartin et al., 2022) to provide useful predictions. At 22°C, we would expect eDNA concentrations to decrease by three orders of magnitude within 96 hours. This decrease, reaching the limit of detection of qPCR assays, is typical of eDNA persistence experiments (McCartin et al., 2022). Hindcasted currents at 60 meters depth at Bright Bank derived from the HYCOM-TSIS 1/100° Gulf of Mexico Reanalysis (Chassignet, 2025) frequently reached and exceeded 0.1 m/s or 0.36 km/hour, a velocity within current speeds measured near other shelf-edge banks in the region (Lugo-Fernández, 1998). Using this value as a static estimate for current speed, and ignoring other processes like diffusion, mixing, and settling, eDNA could be transported away from its source up to 8.64 kilometers per day. Over four days, eDNA may be transported nearly 35 kilometers. Our observations of detectable eDNA ∼1.5 km from its likely source are more than an order of magnitude smaller than this theoretical upper limit.

### 4.3 PCR primer and protocol choice influences eDNA spatial resolution

In our study we compared results using PCR primers and a library preparation protocol designed to broadly amplify marine invertebrates (18S) to PCR primers and a protocol designed to be taxonomically specific to corals (28S). We found that sequencing using the taxonomically specific 28S primers consistently detected coral eDNA at greater distances away from Bright Bank than using the *18S* primers. This difference likely arises from (1) the low relative abundance of coral eDNA relative to other taxa and (2) amplification biases inherent in mixed-template PCR reactions (Kelly et al., 2019). In this case, more abundant sequences (e.g. from copepods) could have been favored over coral sequences using the *18S* primers.

Using taxonomically specific primers improves detection sensitivity and taxonomic classification for their target taxa; however, using specific primers and protocols with sensitive detection thresholds may risk extending detection ranges beyond their immediate sources. Thus, primer specificity and protocol sensitivity critically shape the interpreted spatial distribution of eDNA using metabarcoding and may influence marine biodiversity assessment outcomes with management implications. The effect we observed likely extends beyond the 28S primers used in this study to other taxonomically specific metabarcoding primers (e.g. MiFish) (Miya et al. 2015) and species-specific qPCR and ddPCR assays. Failure to account for this effect could impact the spatial accuracy of biodiversity assessments and species detections using eDNA. Conversely, using primers that broadly amplify many taxa will deliver community-level insights but at reduced sensitivity for ecologically important taxa that are less abundant or shed less eDNA, such as corals. These results underscore the importance of explicitly incorporating both ecological and methodological factors into marine biodiversity monitoring strategies, ensuring accurate and effective spatial management and surveillance practices using eDNA. Empirically comparing the sensitivity of different primer sets on samples with known proportions of DNA from different taxa (i.e. mock communities) can help the assessment of these biases (e.g. Macher et al., 2023).

### 4.4 Study Limitations and Future Directions

This study provides essential insights into eDNA dispersion dynamics but also highlights several current limitations. The Gulf of Mexico is an oceanographically complex region with seasonal variation (Lugo-Fernández, 1998; Bracco et al., 2019). Our empirical findings provide valuable baseline information for interpreting spatial scales of eDNA signals. Dispersal patterns are likely depth-dependent and vary seasonally with changes in oceanographic conditions, and studies investigating seasonal variability in eDNA dispersal are needed. Sampling during one season restricts the temporal generalizability of dispersal distances that can be yielded from our data and similar studies. Seasonal variability in stratification, current strength, and biological processes likely influence eDNA dispersion, warranting longitudinal studies across different seasons and environmental conditions.

Secondly, metabarcoding approaches are semi-quantitative, limiting precise quantification of absolute eDNA concentrations. Integrating metabarcoding with multiple primer sets—including both broad-range and taxon-specific markers—and quantitative methods (qPCR/ddPCR) will offer the most robust strategy for environmental monitoring, spatial biodiversity assessments, and ecosystem management decisions.

Finally, explicit high-resolution hydrodynamic modeling (100 m or less) incorporating physical oceanographic data (e.g., current speed and direction) and field eDNA measurements would significantly refine interpretations of observed dispersal distances. Future research should integrate these empirical and modeling approaches for improved spatial accuracy in marine eDNA studies.

## Author contributions

SH & JMM conceptualized and designed the study. SH, JMM, AFG & LJM acquired the field samples. SH, JMM and AFG acquired funding. LJM performed laboratory work and data analysis and drafted the manuscript. All authors contributed to the interpretation of the data and the writing.

## Data Archiving Statement

18S and 28S metabarcoding data generated in this study are available at the NCBI Sequence Read Archive (SRA) in BioProject PRJNA1159220. Code and necessary data for data analysis are available on FigShare at 10.6084/m9.figshare.28761185.

## Acknowledgements

This research was funded by the National Oceanic and Atmospheric Administration’s Oceanic and Atmospheric Research Office of Ocean Exploration and Research, under award NA18OAR0110289 to SH and JMM at Lehigh University and by NOAA’s National Centers for Coastal Ocean Science, Competitive Research Program under award NA18NOS4780166 to Santiago Herrera at Lehigh University. SH was supported by NOAA award NA18OAR0110289 and the National Academies of Sciences, Engineering, and Medicine Gulf Research Program Early-Career Fellowship under award 2000013668. This research is part of the Woods Hole Oceanographic Institution’s Ocean Twilight Zone Project, funded as part of The Audacious Project housed at TED. This work was also supported by a Lehigh University CAS Dean’s Opportunity Grant.

Special thanks to the Flower Garden Banks National Marine Sanctuary staff (G.P. Schmall, Emma Hickerson, Marissa Nuttall, and Michelle Johnston) for their support of this study. We thank the expedition science team: Dr. Dana Yoerger, Justin Fujii, Eric Glidden, Rene Francolini, and Katie Foley, as well as the captain and crew of *R/V Manta*. We thank Jimmy MacMillan for his assistance with Niskin bottle rosette deployments. We thank Rene Francolini for her contributions to eDNA extraction. We thank Weihua Wang from the Genomics Research Core at the University of Illinois Chicago, and Dr. Stefan Green, Ashley Wu, and Cecilia Chau from the Genomics and Microbiome Core Facility at Rush Medical University for library preparation and sequencing. We thank Dr. Wynn Meyer for her helpful comments on a draft of this manuscript. We also thank Dr. Allen Collins and Annemarie Wood for their helpful discussion regarding interpreting the zooplankton detections in the 28S data.

## Conflict of Interest Disclosure

The authors declare no conflicts of interest.

## Notes

https://doi.org/10.6084/m9.figshare.28761185.v1

### Competing Interest Statement

The authors have declared no competing interest.

### Summary of Updates

This version of the manuscript has been revised to be more concise. All elements of the manuscript (text, supporting information, and figures) have been updated.

