## Appendix A: Supplementary Methods for "Limited Dispersal of Benthic Environmental DNA from a Subtropical Mesophotic Shelf-Edge Bank"

*Detailed Bioinformatic Processing*

Primer sequences were trimmed from forward and reverse sequencing reads using *cutadapt* (Martin, 2011) with the following settings: forward and reverse primer sequences were anchored to the beginning of the forward and reverse reads, respectively; reverse complements of the reverse and forward primers were optionally trimmed from the forward and reverse reads, respectively, if they were identified; the minimum overlap length was set to five nucleotides (-O 5); the number of primer sequences matched was set to one (-n 1); the minimum length of trimmed reads retained was set to five nucleotides (--minimum-length 5); and untrimmed read pairs were discarded if either the forward or reverse primer sequences were not matched and trimmed from the forward and reverse reads, respectively. Read pairs were quality filtered, denoised to infer error-corrected amplicon sequence variants (ASVs), and merged using the *R* package *DADA2* (Callahan et al., 2016). *18S* and *28S* read pairs were filtered using the *filterAndTrim* function with a maximum of two expected errors allowed in either the forward or reverse read. For *18S*, the forward and reverse reads were truncated to 125 bp, and for *28S*, the forward and reverse reads were truncated to 250 and 200 bp, respectively. The pseudo-pool option was used for denoising with the *dada* function (pool = "pseudo"). Chimeras were removed from the resulting ASV tables using the *removeBimeraDenovo* function and the default settings. For the *18S* data, each sequencing run was processed separately, and then ASV tables were merged using the *gather* and *summarise* functions from the *tidyverse* (Wickham et al., 2019).

Taxonomic classification of ASVs was conducted using the *assignTaxonomy* function in *DADA2*, which implements a naïve Bayesian classifier (Wang et al., 2007). Custom reference database files were created from *18S* and *28S* data downloaded from MetaZooGene (https://metazoogene.org/database/) (O'Brien et al., 2024). For *18S*, global data for all taxa were downloaded to form the reference database on April 25, 2024. For *28S*, data from genus and/or species known from the North Atlantic were downloaded for all taxa other than anthozoan corals (Orders: Antipatharia, Scleractinia, Malacalcyonacea, and Scleralacyonacea) on June 20, 2024. A separate, custom reference database with sequences from these anthozoan coral taxa and their most up-to-date taxonomic nomenclature (McCartin et al., 2024) was concatenated to the *28S* data from MetaZooGene to build a comprehensive taxonomic reference database for all animals. For taxonomic classification, the default settings were used, which includes a minimum bootstrap cutoff score of 50. Following taxonomic classification, separate *phyloseq* (McMurdie and Holmes, 2013) objects were created from the ASV tables, taxonomic classification tables, and sampling metadata files for the *18S* and *28S* data. The resulting *phyloseq* ASV tables were "decontaminated" of ASVs that were more prevalent in the negative control samples (both sampling and PCR negative controls) than in the field samples using the *R* package *decontam* and the "prevalence" method with the default settings (Davis et al., 2018).

*Correction for potential contamination*

In the *18S* data, 117,894 reads of ASVs classified to animal phyla were produced in a library prepared from the sampling negative control corresponding to the Niskin bottle rosette cast 1.0 NM to the east of Bright Bank's center. 116,000 reads of ASVs classified to animal phyla were also produced in a library prepared from a PCR negative control from the same PCR plate as the negative sampling control. Cross- contamination to the same degree was not apparent in the *28S* library prepared from the same sampling negative control (602 reads), nor from the

*18S* library prepared from the other sampling negative control on the same PCR plate and just one column away (7 reads). The maximum number of reads in any of the other sampling negative controls or PCR negative controls in the *18S* dataset was 334 reads. We assumed that cross contamination of the negative sampling control corresponding to the cast east of Bright Bank occurred during *18S* library preparation, and we excluded all samples from this cast for the analysis of the *18S* data out of caution.

In the *28S* data, >100 sequencing reads were produced from libraries prepared from several sampling negative controls. We detected 279 total sequencing reads in the sampling negative control corresponding to the cast 1.5 NM to the NW of Bright; animal ASVs detected in this sample were classified to the black coral genus *Antipathes*. We detected 102 total sequencing reads in the sampling negative control corresponding to the cast 3.0 NM to the NW of Bright; animal ASVs detected in this sample were classified to *Antipathes* and the trachymedusae genus *Aglaura*. We also recovered 477, 602, and 5,077 total sequencing reads in the sampling negative controls corresponding to the CTD casts 1.5 NM to the NE, 1.0 NM to the E, and 1.5 NM to the SW of Bright Bank, respectively. Animal ASVs in these samples were classified to the octocoral genera *Muricea* and *Eunicea* (Family: Plexauridae).

We chose to retain sequences from taxa detected in the negative controls for data analysis. The detections of these taxa in field samples could have very likely represented true detections, and false positive detections of these taxa would not have meaningfully influenced our broader interpretation of the data. eDNA sequences from *Muricea* and *Eunicea*, which were most numerous among those taxa detected in the sampling negative controls, were largely constrained to samples from one depth in the CTD cast corresponding to the sampling negative control where they were detected. We infer that the negative control sample was likely contaminated from these few samples that represent valid detections of these genera. *Antipathes* was the most common coral genus across samples, based on both the number of sample replicates in which it was detected (52 of 115) and its average relative abundance across these samples (3.2%). Therefore, its presence in any sample below the surface mixed layer (it was not detected in any sample in the surface mixed layer) was plausible. False positive detections of these three coral taxa in the *28S* data, the trachymedusae *Aglaura* in the *28S* data, and any taxa detected in sampling negative controls at low sequence abundances in the *18S* data (excluding the samples that were removed) may have marginally influenced some analyses that included all the data, for example ordination. However, broader interpretation of patterns in the distribution of eDNA with depth and with distance from the center of Bright Bank would not be changed by their exclusion.
