## Appendix B: Supplementary Figures for "Limited Dispersal of Benthic Environmental DNA from a Subtropical Mesophotic Shelf-Edge Bank"

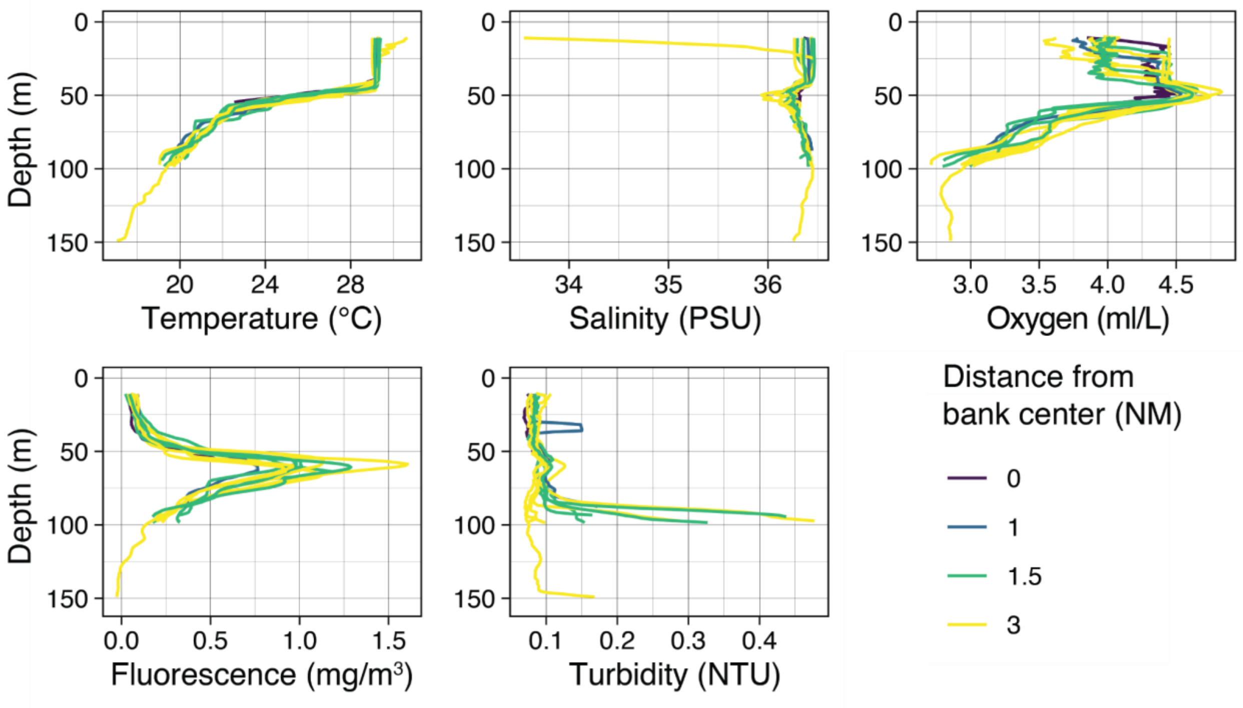


**Figure S1:** Profiles of temperature, salinity, oxygen concentration, fluorescence, and turbidity by depth collected during downcasts of Niskin bottle deployments at and near Bright Bank.

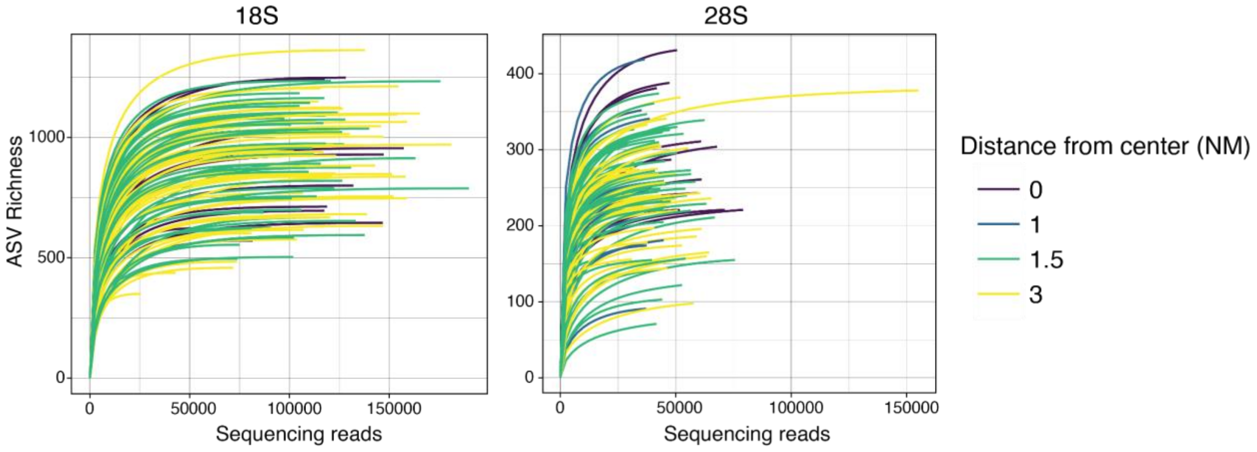


**Figure S2: Rarefaction analysis of eDNA amplicon diversity across samples.** Rarefaction curves showing the accumulation of amplicon sequence variants (ASVs) in increments of 2,500 reads per sample for 18S and 28S markers. Lines represent individual samples. Color indicates the cast’s distance from the center of Bright Bank. Curves illustrate sequencing depth adequacy and richness across sites.

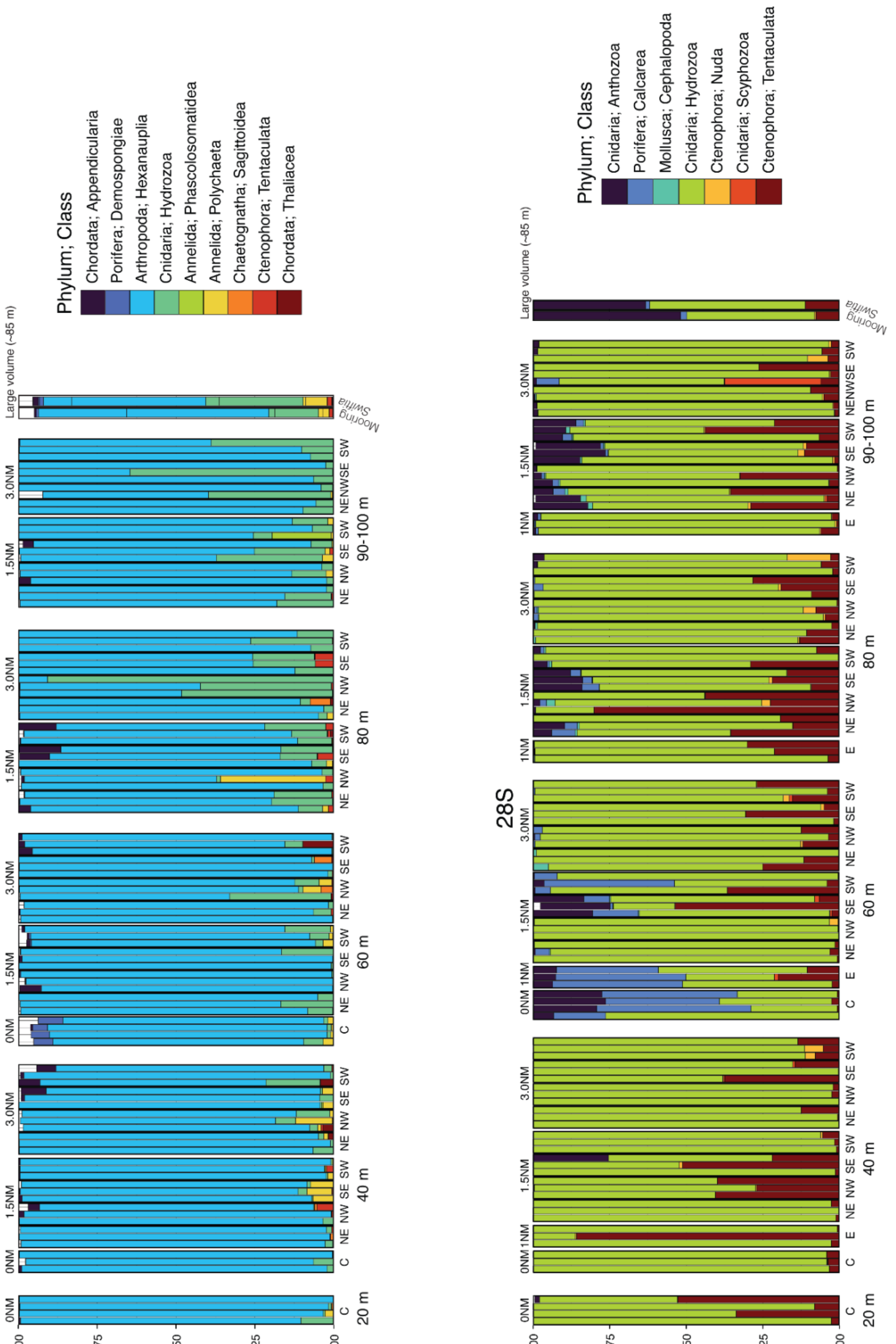

18S
28S
Animal reads
1.00
0.75
0.50
0.25
0.00
0NM
1NM
1.5NM
3.0NM
C
E
NE NW SE SW
20 m
40 m
60 m
80 m
90-100 m
Large volume (~85 m)
Mooring
Swiftia
Phylum; Class
Chordata; Appendicularia
Porifera; Demospongiae
Arthropoda; Hexanauplia
Cnidaria; Hydrozoa
Annelida; Phascolosomatidea
Annelida; Polychaeta
Chaetognatha; Sagittoidea
Ctenophora; Tentaculata
Chordata; Thaliacea
Cnidaria; Anthozoa
Porifera; Calcarea
Mollusca; Cephalopoda
Ctenophora; Nuda
Cnidaria; Scyphozoa

**Figure S3: Proportion of animal eDNA sequencing reads by Class.** Distribution of sequencing read abundances across samples categorized by depth, horizontal distance, and direction from Bright Bank. Directions include: C = Center, E = East, NE = Northeast, NW = Northwest, SE = Southeast, and SW = Southwest.

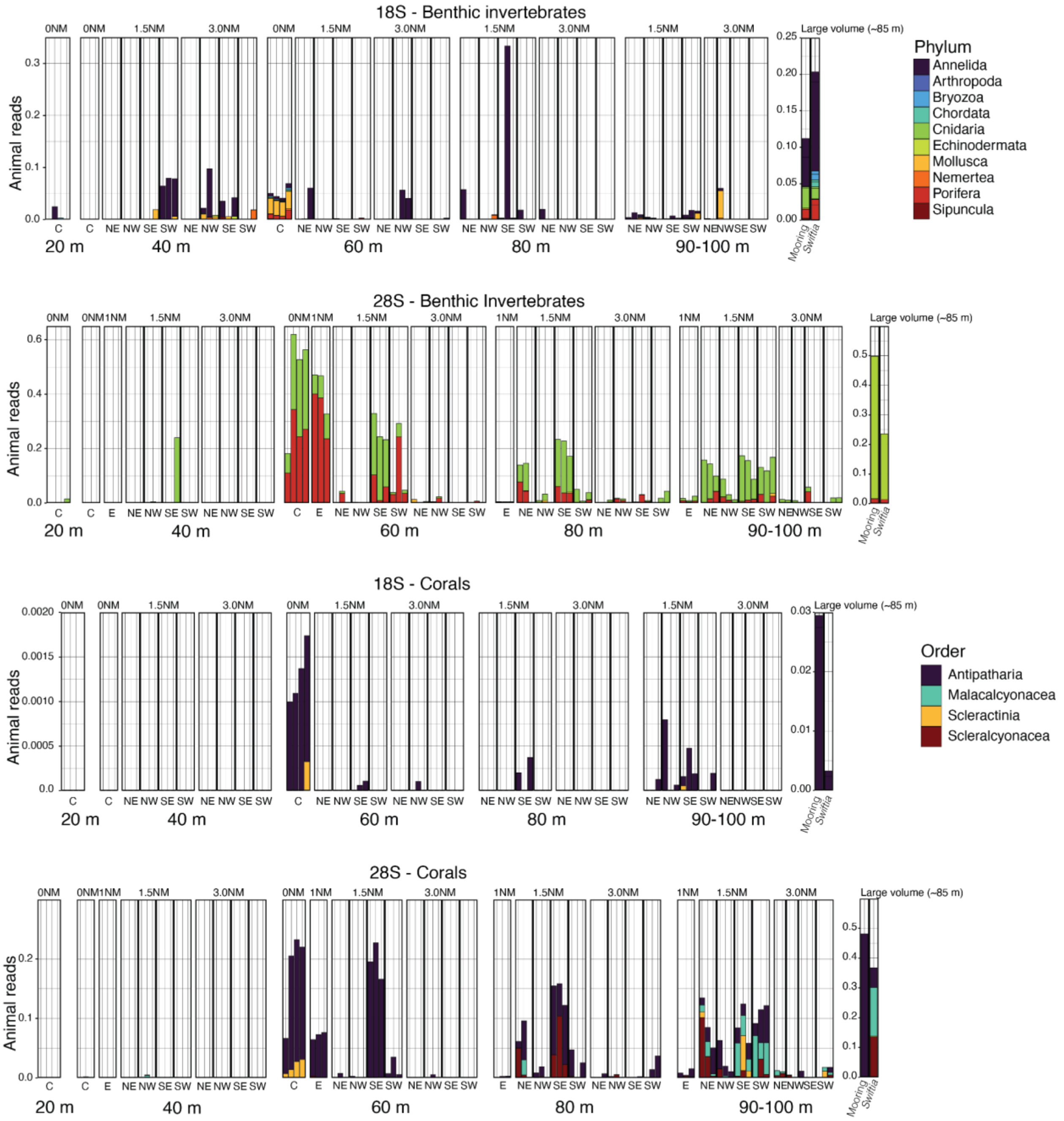

**Figure S4: Proportion of benthic invertebrate eDNA sequencing reads by taxa.**

Bar plots showing proportions of eDNA sequencing reads from benthic invertebrate families across samples arranged by depth, distance, and direction from Bright Bank center. Sample directions abbreviated as: C = Center, E = East, NE = Northeast, NW = Northwest, SE = Southeast, and SW = Southwest. Y-axis scales vary to accommodate sample volume differences.
